# Targeting IRS-1/2 in uveal melanoma inhibits *in vitro* cell growth, survival and migration, and *in vivo* tumor growth

**DOI:** 10.1101/2022.10.26.513928

**Authors:** Chandrani Chattopadhyay, Rajat Bhattacharya, Jason Roszik, Fatima S. Khan, Gabrielle A. Wells, Hugo Villanueva, Yong Qin, Rishav Bhattacharya, Sapna Patel, Elizabeth A. Grimm

**Affiliations:** Departments of Melanoma Medical Oncology, The University of Texas MD Anderson Cancer Center, Houston, Texas-77030, USA; Departments of Genomic Medicine, The University of Texas MD Anderson Cancer Center, Houston, Texas-77030, USA; Departments of Surgical Oncology, The University of Texas MD Anderson Cancer Center, Houston, Texas-77030, USA; Patient Derived Xenograft and Advanced In Vivo Models Core, and Department of Otolaryngology – Head and Neck Surgery, Baylor College of Medicine, Houston, Texas-77030, USA; Department of Pharmaceutical Sciences, The University of Texas at El Paso, El Paso, Texas-79968, USA; College of Agriculture and Life Sciences, Texas A & M at College Station, College Station, Texas-77843, USA

## Abstract

Uveal melanoma (UM) originating in the eye and metastasizing to the liver is associated with poor prognosis and has only one approved therapeutic option. We hypothesized that liver-borne growth factors may contribute to UM growth. Therefore, we investigated the role of insulin-like growth factor −1 and its receptor (IGF-1/IGF-1R) signaling in UM. We found that the insulin receptor substrate −1 (IRS-1) is overexpressed in UM cells and tumors. Since we previously observed that IGF-1R antibody therapy was not clinically effective in UM, we investigated the potential of NT157, a small molecule inhibitor of IRS-1/2 in blocking this pathway in UM. NT157 treatment in UM cells resulted in reduced cell survival and migration, and increased apoptosis. This treatment also significantly inhibited UM tumor growth *in vivo*, in the chicken egg chorioallantoic membrane (CAM) and subcutaneous mouse models, validating the *in vitro* effect. Mechanistically, through reverse phase protein array (RPPA), we identified significant proteomic changes in the PI3K/AKT pathway, a downstream mediator of IGF-1 signaling, with NT157 treatment. Together, these results suggest that NT157 inhibits cell survival, migration *in vitro* and tumor growth *in vivo* via inhibiting IGF-1 signaling in UM.

## Introduction

Uveal melanoma (UM) arises from melanocytes in the choroidal plexus of the eye and is biologically different from cutaneous melanoma due to different genetic alterations [1] and a strong tendency to metastasize to the liver [2, 3] (observed in approximately90% of cases with metastasis). The reason for such preferential liver seeding of UM metastatic cells is unknown. Growth factors have been shown to play a significant role in the tissue-specific homing of cancer cells [4]. Liver-borne growth factors, such as the IGF-1 may contribute to this distant metastasis to liver and blocking these factors might have therapeutic value. IGF-1 plays a major role in tumor cell proliferation and survival, and its receptor, IGF-1R, plays a significant role in multi-step process of cancer metastasis [5]. In UM, IGF-1R expression has been shown to be independently prognostic in multi-variate analysis [6]. Also, IGF-1 is one of the major growth factors secreted from the liver, and high IGF-1 serum levels have been shown to be associated with UM metastasis [7].

Protein expression of IGF-1R, the receptor for IGF-1, has been detected in UM tumors, and its high expression was shown to be associated with poor survival [8–10]. Despite this high IGF-1R expression, few attempts have been made to target this ligand/receptor system in UM. A phase II trial that targeted IGF-1R in UM with a human monoclonal antibody from IMCLONE (NCI #8832, 2010-0451) [11] did not have any therapeutic benefit and concluded that this monoclonal antibody was not a viable treatment option. Targeting the IGF-1 pathway alone or in combination for more effective therapy regimens against metastatic UM remains to be studied [12]. IRS1/2 are the first downstream targets of activated IGF-1R, which transmit IGF-1R mediated signals. The IRS proteins do not have their own kinase activity, but act as scaffolds for signaling complexes to initiate cellular signaling pathways [13]. IRS-1/2 are intermediates of multiple receptors that control tumor progression and thus can play an important role in the response of tumor cells to different microenvironmental signals. NT157, the small molecule IRS1/2 inhibitor, developed by Dr. Alexander Levitzki, functions by serine phosphorylation and destruction of IRS1 and IRS2, leading to long-term blocking of IGF-1R signaling and strong inhibition of tumor growth [14] in ovarian and prostate cancers as well as BRAF inhibitor resistant melanoma.. The NT compounds are highly effective in anti-cancer activity because they inhibit both the tumor growth promoted by IRS1 and the IRS2-mediated metastasis [15]. Furthermore, inhibition of both IRS proteins precludes the compensation of one for the other [14, 15].

In this study, we have utilized both the *in vitro* and *in vivo* models of UM to investigate the effect of IRS-1 inhibition on UM growth, cell survival and migration. Our *in vivo* models, the CAM model and mouse subcutaneous (subQ) xenograft model [16], were previously tested and standardized for studying UM growth inhibition [17]. Here, we show, *in vitro* treatment with NT157 leads to inhibition of UM cell growth, induction of apoptosis and inhibition of cell migration. More significantly, we show inhibition of UM tumor growth on CAM as well as in our subQ mouse model with NT157 treatment.

## Materials and Methods

### Antibodies and reagents

The IRS-1 antibody for immunohistochemistry analysis was obtained from ABCAM (#ab40777; Abcam, Boston, MA). Similarly, Caspase 3 antibody was obtained from ABCAM (#ab184787). Cell signaling Danvers, MA provided antibodies for Caspase 9 (#9502S); Hexokinase (#2867S), STAT3 (#4904S), Phospho STAT 3, (#9131S), Cleaved PARP (#5625S), IRS-1 for western blotting (#2382S), Phospho p38 MAP Kinase (#9215S), p38 MAP Kinase (#9212S), Phospho AKT (#3787S), AKT (#9272S), Phospho Erk1/2(#4376S), Erk1/2 (#9102S) and IGF-1R (#3027S). Phospho IGF-1R antibody was obtained from Sigma Aldrich (SAB4300652; Millipore Sigma, Burlington, MA). The Actin antibody was obtained from Santa Cruz (#sc-47778; Santa Cruz, Dallas, TX). NT157 was obtained from Selleckchem (#NM-4126; Houston, TX).

Cell migration assay plates were obtained from Corning (#354578; Corning, Glendale, AZ) and staining Kit was obtained from Siemens (#B4132-1A; Siemens Health care Diagnostic Inc, Newark, DE). MTT reagent was obtained from EMD Millipore Corp (#475989; United Kingdom). Propidium Iodide was procured from Sigma-Aldrich (#81845; Millipore Sigma, Burlington, MA). Ribonuclease A (RNase) was obtained from Sigma-Aldrich (#R4642; Millipore Sigma, Burlington, MA).

### Cell culture and treatments

Cell line Mel20–06–039, (RRID:CVCL_8473) [18, 19], was obtained from Dr. Tara A. McCannel. Cell line OMM-1 (RRID:CVCL_6939) [12], Mel202 (RRID: CVCL_C301), 92–1 (RRID: CVCL_8607) [12], and Mel270 (RRID: CVCL_C302) [12] were kindly provided by Drs. Martine Jager and Bruce Ksander. Cell lines obtained from ATCC were: MM28, MP38, MP41, MP46, MP65. Additional UM cell line information is provided in the references [20–23].

UM cells were cultured in RPMI 1640 media with 10% FBS, glutamine, Penicillin-streptomycin and Insulin supplement, under ambient oxygen at 37 °C.

### Cell line validation

Cell lines were validated by short random repeat (STR) DNA fingerprinting techniques and mutational analysis, by the MDACC Cancer Center Support Grant (CCSG)-supported Characterized Cell Line Core, using the AmpFLSTR Identifier Kit (Applied Biosystems, Foster City, CA), according to manufacturer’s instructions. The STR profiles were compared to known ATCC fingerprints (http://www.ATCC.org), and to the Cell Line Integrated Molecular Authentication database (CLIMA) version 0.1.200808 (http://bioinformatics.istge.it/clima/). The STR profiles matched known DNA fingerprints or were unique.

### Western blotting

Cells were lysed in a buffer containing 50 mM Tris (pH 7.9), 150 mM NaCl, 1% NP40, 1 mM EDTA, 10% glycerol, 1 mM sodium vanadate and a protease inhibitor cocktail (Roche Pharmaceuticals, Nutley, NJ). Proteins were separated by SDS-PAGE with 4-20% gradient gels (Bio-Rad Laboratories, Hercules, CA), transferred to a Hybond-ECL nitrocellulose membrane (GE Healthcare Biosciences, Piscataway, NJ) and blocked in 5% dry milk in PBS. The membrane was then incubated with primary and secondary antibodies, and target proteins were detected with ECL detection reagent (GE Healthcare Biosciences).

### FACS Flow cytometry analysis for cell surface expression of IGF-1R

UM cells were washed with phosphate-buffered saline (PBS) and detached by gentle pipetting. The cells were incubated with PE-conjugated mouse monoclonal anti-human IGF-1R antibody (BD Biosciences, Franklin Lakes, NJ) or PE-conjugated isotype-matched control mouse antibody (IgG1k, BD Biosciences) for 1 h on ice and then washed with PBS. The cells were analyzed using FACSCalibur (BD Biosciences), and the data were analyzed using FlowJo software (Tree Star, Inc.).

### Propidium Iodide staining and flow cytometry analyses

UM cells were trypsinized, washed and fixed with 70% Ethanol and kept at 4 °C overnight. Cells were centrifuged and pellet resuspended in PBS to rehydrate for 15 min. Cells were then centrifuged at 500 g and treated with 200 μg/mL RNase A for 1 h at 37 °C. Cells were stained with 40 μg/mL Propidium Iodide for 20 min at room temperature. After centrifugation at 500 x g, cells were resuspended in PBS with 0.02% EDTA for cell cycle analysis using a Beckman Coulter Galios 561 analyzer.

### Colony formation assay

For the colony formation assay, UM cells were seeded in 24 well plates at 500 cells/well. Next day NT157 (1 and 2.5 μM) was added to the wells. Controls were untreated cells. Cell culture media and NT157 were replaced every 3 days. Colonies were allowed to grow until clones in the control wells covered 70-80% of the well surface. For fixing and staining the cells, the culture media was removed, wells washed with 1X PBS to remove dead cells and 1ml of crystal violet in 25% methanol was added. Cells were stained for 5 min at RT. Crystal violet was aspirated and the wells washed with water until crystal violet was removed from everywhere else but the colonies.

### Reverse phase protein array (RPPA) analysis

RPPA analyses were performed at the UT MD Anderson Cancer Center’s Functional Proteomics RPPA Core facility. Briefly, cell lysates were two-fold serially diluted for five dilutions (from undiluted to 1:16 dilution) and arrayed on nitrocellulose-coated slides. Samples were probed with antibodies using catalyzed signal amplification and visualized by 3,3’-diaminobenzidine colorimetric reaction. Slides were scanned on a flatbed scanner to produce 16-bit TIFF images of the reacting spots, and spot densities were quantified using the MicroVigene software program. Relative protein levels for each sample were determined by interpolation of each dilution curves from the “standard curve” constructed by a script in R written by MD Anderson’s Department of Bioinformatics & Computational Biology. Heatmaps were generated in Cluster 3.0 (http://www.eisenlab.org/eisen/) as a hierarchical cluster using Pearson correlation and a center metric.

### Chicken egg chorioallantoic membrane (CAM) tumor xenograft model

UM cell line 92.1 engineered to express luciferase was engrafted on the CAM for 7 days following previously established methods [24, 25]. Briefly, embryonic day 7 eggs were inoculated with 5 × 10^5^ cells in a 1:1 Matrigel and PBS (supplemented with calcium and magnesium) solution. Eggs were randomized into 2 groups with 12 eggs in each set. Three days later, daily vehicle and NT157 (1 μM) treatments were topically applied. Bioluminescence imaging was performed at days 5 and 7 post engraftment using an IVIS Lumina III in vivo imaging system (Perkin Elmer, Waltham, MA). IVIS instrument exposure time was 3 seconds, 15 minutes after addition of 15 mg/ml D-luciferin. Region of interest and total flux was determined using the Living Image Software suite. Tumors were imaged and harvested at day 7 post engraftment. Chick embryos were euthanized per AVMA guidelines. CAM tumors were fixed in 10% formalin and embedded in paraffin blocks for downstream immunohistochemical analysis. IHC was performed on paraffin sections of CAM tumors using a pan melanoma antibody cocktail (Biocare Medical, Pacheco, CA).

### Cell viability assays

Methylthiazole tetrazolium (MTT)- based cell viability assays were used for estimating cell survival. UM cells were plated at a density of 1 × 10^4^ cells/well in triplicate in a 24-well plate. To assess cell viability, MTT reagent [3-(4,5-dimethylthiazol-2-yl)-2,5-diphenyltetrazolium bromide] (Sigma-Aldrich, St. Louis, MO), dissolved in PBS, was added to a final concentration of 1 mg/mL. After 3 h the precipitate formed was dissolved in DMSO, and the color intensity estimated in a MRX Revelation microplate absorbance reader (Dynex Technologies, Chantilly, VA) at 570 nm.

### Cell migration assay

Cell migration assays were performed in Boyden chambers using uncoated filters (BD Biocoat control inserts, BD Biocoat, San Jose, CA). UM cells were plated in 10 cm dishes and treated overnight with NT157 (2.5 μM) or MAB391 (10 μg/ml) for 48 h. Untreated cells were used as control. Next day, 1×10^5^ cells/well were plated in serum-free medium, and the migration assay done with 10% FBS as chemo-attractant and procedure as described in Chattopadhyay et al., [26]. Stained cells were photographed with a Nikon Eclipse TE2000-U microscope at 20X magnification using NIS Elements advanced research software. To quantify migration, the cells in each filter were counted from five independent fields under the microscope at 40X magnification and the mean cell number/field was calculated. Each assay condition was tested in 2 replicates.

### NT157 treatment of subQ UM tumors in NSG mice

For *in vivo* studies UM cell lines, 92.1 and MM28, were grown subcutaneously in NOD *scid* gamma mice. 0.5 × 10^6^ UM cells were suspended in 50 μl of HBSS media + growth factor reduced Matrigel (1:1 ratio) and subcutaneously injected into the right flank of each mouse. Tumors were measured by slide calipers and allowed to grow to ~75-100 mm^3^. The animals were then randomized for treatment with vehicle (40% PEG300 + 5% Tween 80 in water) or NT157 (50 mg/Kg), with 5 animals for each treatment set. Drugs were injected intraperitoneally on alternate days for a total of 3 days each week till end of experiment. Tumor sizes were measured twice every week using slide calipers by blinded observers. All *in vivo* studies were performed in accordance with accepted guidelines for housing, euthanasia and treatment, under an Institutional Animal Care and Use Committee-approved protocol at MD Anderson.

## Results

### IGF-1R and IRS-1 are expressed in uveal melanoma

Since IGF-1 is expressed in highest saturation in the liver [27, 28], and liver is the primary site for UM metastasis [29] we checked the expression of corresponding receptor IGF-1R and the immediate substrate of the receptor activation, IRS-1, in our UM cells and primary tumors. We analyzed 31 unique tumor types and their corresponding normal tissues, including UM tumors from The Cancer Genome Atlas (TCGA) data for IGF-1R gene expression. Each dot on the IGF-1R expression plot (Figure 1A) represents one tumor (red) or normal (green) sample and the samples above the median (expression line) are considered to be overexpressed compared to their corresponding normal tissues. Also, the IGF-1R expression levels are progressively higher moving from left to the right most end of the plot. UM tumors displayed the third highest IGF-1R gene expression amongst cancer types, surpassed only by prostate adenocarcinoma (PRAD) and breast invasive carcinoma (BRCA) (Figure 1A). Next, we screened for IGF-1R RNA and protein expression in validated human UM cell lines representing known genetic backgrounds. All lines showed IGF-1R gene expression as indicated by the RT PCR analysis (Figure 1B), and phosphorylation of IGF-1R was found to be inducible by the ligand, IGF-1, treatment, demonstrating it can be activated in most lines (Figure 1C). Higher expression of IGF-1R was observed in UM cells as compared to non-cancer cell lines (FMC15H melanocytes, HaCaT keratinocytes and BJ fibroblasts) (Figure 1D). The cell surface expression of IGF-1R protein on UM cells was detected by flow cytometry analysis (Figure 1E). In order to confirm the expression of downstream pathway components in UM, we investigated the expression of IRS-1, the immediate downstream effector of IGF-1R. Using TCGA data sets, we found IRS-1 transcript expression in UM human patient samples (Figure 2A). Immunohistochemical analysis of IRS-1 protein expression in a matched set of primary and metastatic UM tissues, from the eye and the liver, are shown in Figure 2B. Therefore, UM cells express both IGF-1R and IRS-1 in detectable levels.

**Figure 1.**
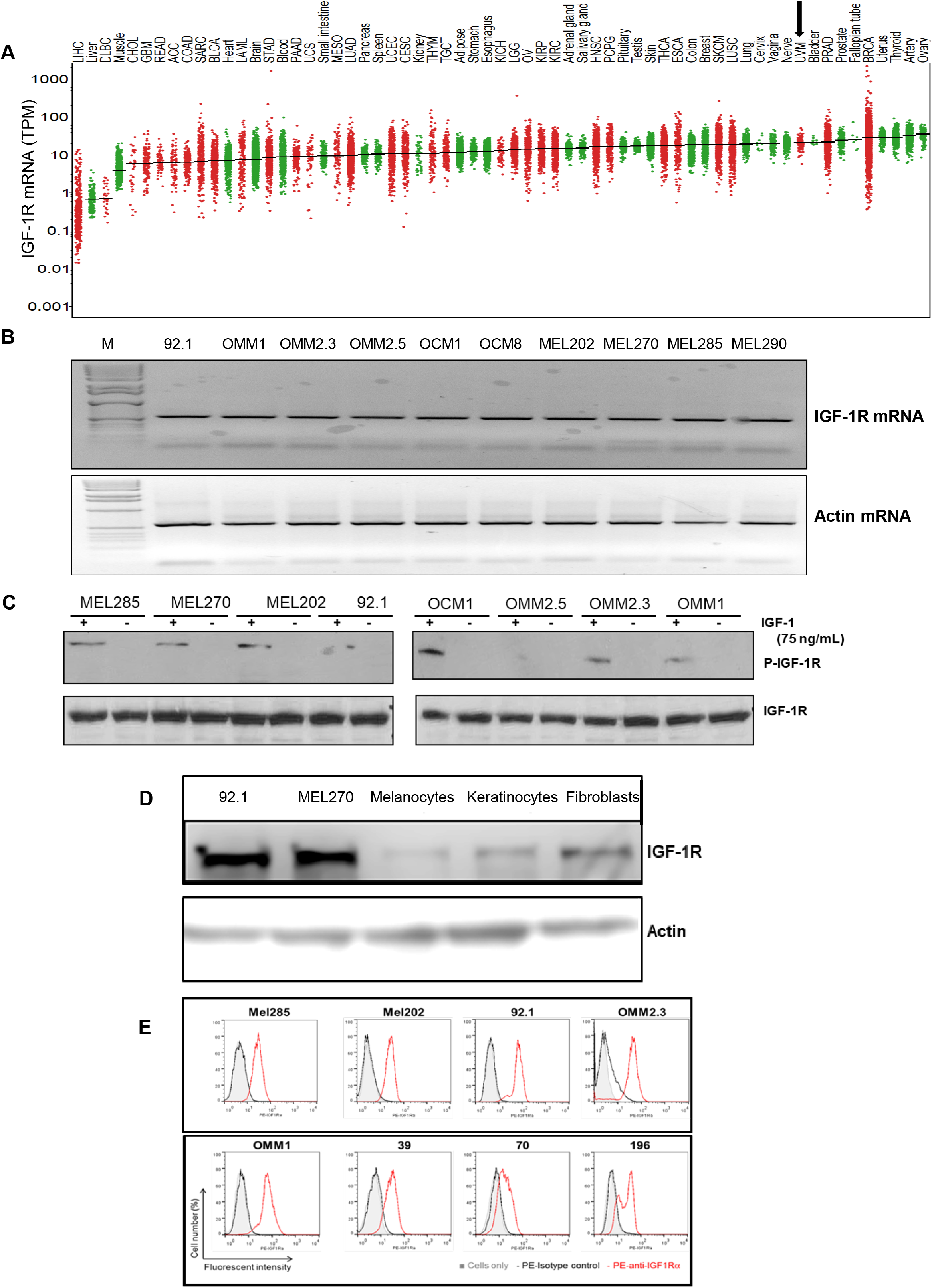
Expression of IGF-1R in UM tumors and cell lines. (**A**) The IGF-1R mRNA expression levels in tumor and normal tissues were compared in 33 cancer (red dots) and corresponding normal (green dots) tissues through TCGA database analysis. The expression in UM is indicated by an arrow. (**B**) RT-PCR analyses showing IGF-1R mRNA levels in 10 different UM cell lines. (**C**) Western blot analysis showing IGF-1R activation/induction in UM cell lines with 75 ng/mL of IGF-1 treatment. (**D**) Western blot analysis for IGF-1R levels using IGF-1R antibody shows higher expression of IGF-1R protein in UM cell lines (92.1 and Mel270) *versus* non-cancerous melanocytes, keratinocytes and fibroblasts. (**E**) FACS histogram plots showing cell surface expression of IGF-1R in the UM cell lines (Mel285), Mel202, 92.1, OMM2.3, OMM1, 39, 70 and 196) stained with IGF-1R antibody (red lines). The isotype control is represented by the black lines in the FACS histograms.

**Figure 2.**
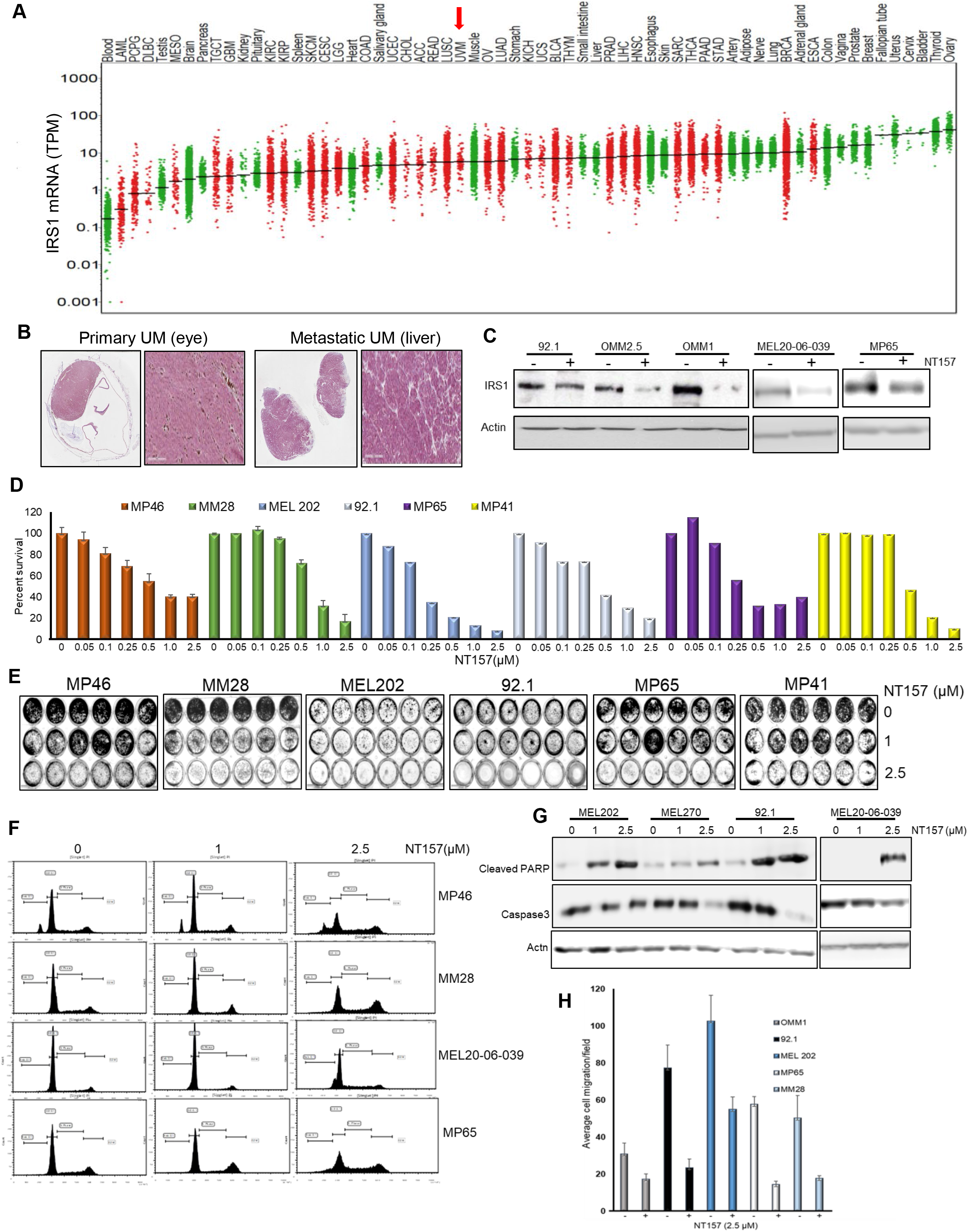
NT157 treatment reduces IRS-1 levels leading to reduction of cell viability, migration and induction of apoptosis in UM cell lines. (**A**) TCGA dataset analysis shows high expression of IRS-1 mRNA in UM tumors (indicated by an arrow) among 33 cancer (red dots) and corresponding normal (green dots) tissues. (**B**) Immunohistochemical staining of matched primary (eye) and metastatic (liver) UM tumor tissues using IRS-1 antibody detects IRS-1 expression in both eye and liver. (**C**) Western blot analysis of cell extracts from UM cell lines (92.1, OMM2.5, OMM1, MEL20-06-039, MP65) treated with NT157 (1 μM) and probed with anti-IRS-1 antibody shows IRS-1 protein levels decrease in UM cell lines with NT157 treatment. (**D**) Quantification of percent cell survival using MTT-based assays in UM cell lines (MP46, MM28, MEL202, 92.1, MP65 and MP41) with (0.05, 0.1, 0.25, 0.5, 1.0, 2.5 μM) or without NT157 treatment shows NT157 dose-dependent decrease in cell survival. The bar graphs represent percentage of cell survival and are a mean ± SD of three independent experiments. (**E**) **Colony formation assay**: Representative images of crystal violet staining shows fewer number of colonies formed by UM cell lines (MP46, MM28, MEL202, 92.1, MP65 and MP41) with NT157 treatment (1 and 2.5 μM) compared to untreated controls. (**F**) Representative FACS histograms of DNA content detected by propidium iodide staining to detect cell cycle status in UM cell lines (MP46, MM28 and MEL20-06-039 and MP65) shows a dose-dependent increase of number of cells in G0/G1 phase with NT157 treatment (1 and 2.5 μM) *vs*. no treatment. (**G**) Western blot analysis of total cell extracts shows NT157 dose-dependent (1 and 2.5 μM) increase in cleaved PARP and decrease in caspase3 levels detected by respective antibodies in UM cell lines (MEL202, MEL270, 92.1 and MEL20-06-039) compared to untreated controls. (**H**) Quantification of cell migration in UM cell lines (OMM1, 92.1, MEL202, MP65 and MM28) shows reduction in migration of cells treated with NT157 (2.5 μM) compared to untreated controls. An average of five fields of cells/filter were counted under a microscope with 40X magnification. The average number of cells counted/field were obtained in two independent experiments and the mean ± SD of these average cell counts/ field were plotted as bar graphs.

### IRS-1 inhibiton reduces survival, induces apoptosis, and inhibits migration of UM cells

Small molecule inhibitor NT157 inhibits its target, IRS-1, by protein degradation [14]. We treated UM cells expressing IRS-1 with NT157 to inhbit IRS-1 activity, and observed a reduction in IRS-1 protein levels as shown in western blots (Figure 2C). Since reduction in IRS-1 is expected to affect cell survival [15] so we tested the effect of NT157 on UM cell survival. Through MTT assays we observed inhibition of cell survival in all cell lines tested (Figure 2D). UM cell lines varied in sensitivity to NT157 treatment (2.5 μM), ranging between 40%-85% reduction in cell survival. We also corroborated the above observation through the colony formation assay. We treated MEL202, 92.1, MM28, MP38, MP41, MP46, MP65 UM cell lines with NT157 in the colony formation assay, and observed inhibition of colony growth or reduction in colony number (Figures 2E and S2). We again observed varied sensitivities to NT157 among the cell lines tested.

We also observed that NT157 induces apoptosis in UM cells, as evident from dose-dependent increase in sub G1 population upon NT157 treatment, observed through FACS analysis after propidium idodide staining (Figure 2F and Table 1). To charecterize whether this G0/G1 cell accumulation is due to apoptosis, we performed molecular analysis of apoptotic marker using western blotting. We observed increased levels of PARP processing (cleaved PARP induction) as well as caspase 3 processing (reduction in caspase 3 protein levels) after NT157 treatment of UM cell lines (western blots, Figure 2G).

**Table 1.** Shows percent cells gated in subG1 phase with /without treatment with NT157. The percent of cells in cell cycle phases were obtained by counting propidium iodide stained cells after 48 h of NT157 treatment. At the higher NT157 concentration we see an accumulation of subG1 cells compared to the untreated controls.

As IGF-1 pathway is a known modulator of cell migration [30], we tested the effect of IRS-1 inhibiton on UM cell migration. The UM cell lines treated with NT157 overnight were subjected to a Boyden Chamber-type cell migration assay with 10% FBS as the chemo-attractant. Significantinhibition of cell migration was observed in NT157-treated UM cells (Figures 2H and S1B).

We also compared the efficacy of NT157 in its ability to block different cellular/physiological processes against a neutralizing, monoclonal antibody to IGF-1R (MAB391). After confirming MAB391 targets IGF-1R activation (Figure S1A), we compared the effect of MAB391 vs. NT157 in cell viablity and migration assays. NT157 was more effective in blocking UM cell survival in an MTT-based assay in the two cell lines tested (Figure S1B) when compared with MAB391. Similarly, in an *in vitro* migration assay, NT157 performed better in cell migration inhibition in response to 10% fetal bovine serum (Figure S1C). Overall, NT157 blocks the physiological functions of IGF-1/IGF-1R pathway in UM cells with high efficiency.

### Multiple cellular pathways are modulated in response to IRS-1 inhibition

Since IGF-1R affects several different signaling pathways in the cells, we wanted to understand how the major cellular signaling pathways in UM cells will be affected by NT157 treatment. Proteomic analysis with Reverse Phase Protein Array (RPPA) was performed after treatment of four different UM cell lines with two NT157 doses (1 μM and 2.5 μM) and for two different time intervals (24 and 48 h). RPPA data analysis of NT157-treated vs. untreated UM cells demonstrated several changes in major cell signaling pathways. We observed dose- and time-dependent increases in p85, the inhibitory subunit of PI3K catalytic activity compared to untreated samples, which indicates reduced activation of PI3K pathway. We also observed minimal increase in the activation of MAPK pathways, and changes in cell cycle effectors and increase in hexokinase-II levels as shown in Figure 3A. To validate the signaling pathway changes from RPPA results, we used wesern blotting on an independent set of samples to detect significant changes observed. Hexokinase-II seemed to increase in an NT157 dose-dependent manner in all cell lines tested. Similarly, JNK unformly decreased with increase in NT157 dose. At higher concentration (2.5 μM) and longer timepoint (48 h) AKT activation was effectively reduced (Figure 3B). Therefore, this data shows a reduction in overall activated levels of PI3K/AKT pathway upon NT157 treatment, indicating an inhibitory effect of NT157 on UM cell survival.

**Figure 3.**
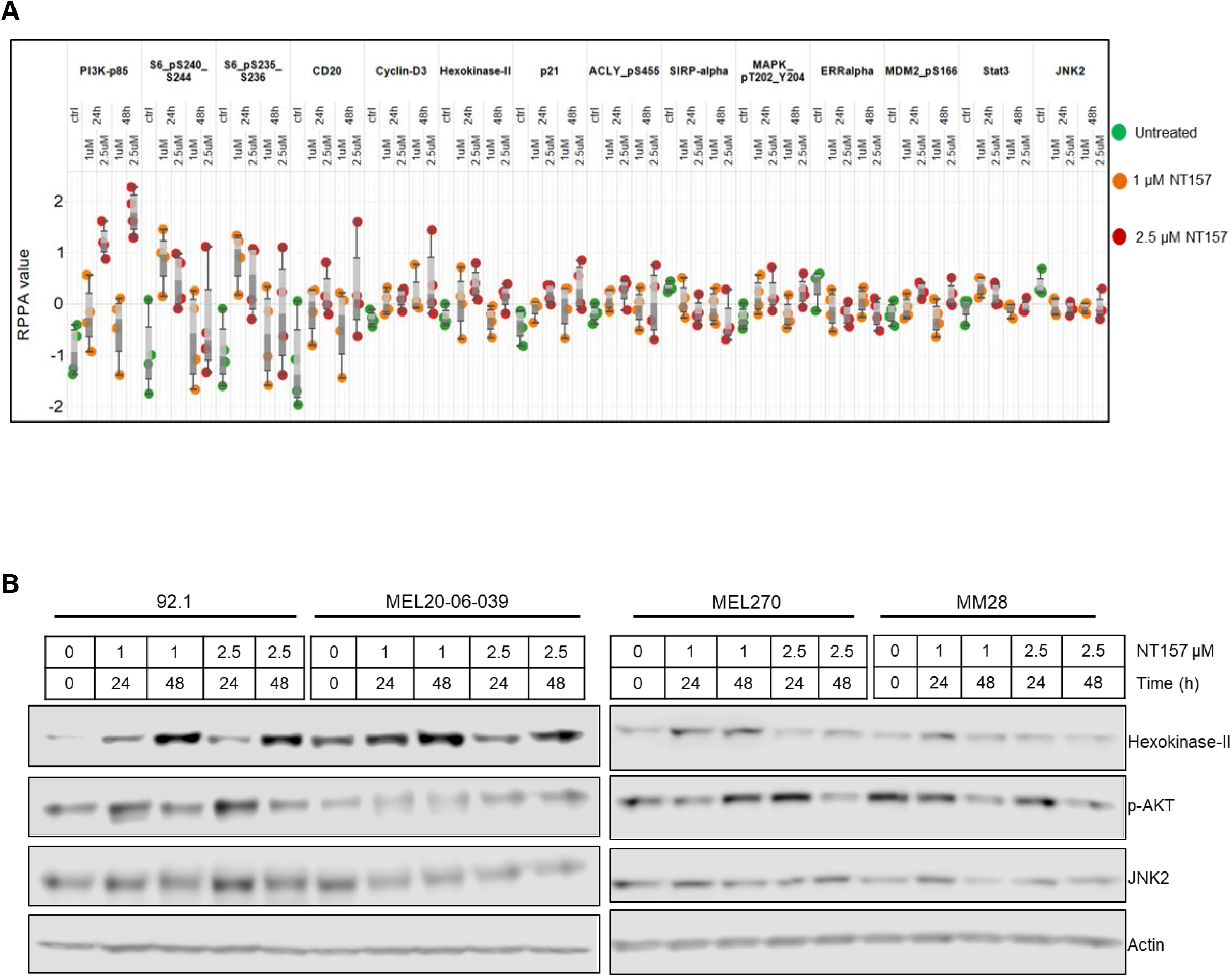
Significant changes in cell signaling proteins detected post-NT157 treatment. (**A**) RPPA protein profiling of four 4 UM cell lines (MEL270, MM28, MEL-20-06-039 and 92.1; represented by each dot). The RPPA expression values are displayed on the y-axis for antibodies that show a significant difference (*P* < 0.05) compared to control sample. Cells were treated either for 24 or 48 hours with a 1 or 2.5 μM concentration of NT157. Green dots represent untreated cells; orange and red dots represent cells treated with 1 and 2.5 μM of NT157, respectively for 24 and 48 h. (**B**) Western blot analyses to validate some of the significant changes observed in RPPA using antibodies against hexokinase-II, phospho-AKT, and JNK2 shows treatment and time-dependent (1 and 2.5 μM for 24 and 48 h) upregulation of hexokinase-II and downregulation of phospho-AKT.

### IRS-1 targeting reduces UM tumor growth in a chicken CAM model

The chicken CAM has been previously used as a successful *in vivo* system to model UM xenograft growth [17, 31]. To test the effect of IRS-1 inhibition on tumor progression *in vivo*, we engrafted luciferase-tagged 92.1 cells (a UM cell line) on the chicken CAM to generate UM xenografts. After confirming that the tumors generated on the CAM model are of melanoma origin by pan-melanoma marker staining (HMB1, Tyrosinase and S100) (Figure 4A), we treated the resulting tumors with different concentrations of NT157. Gross tumor analysis demonstrates a reduction in overall tumor growth in response to NT157 treatment (Figure 4B).The bioluminescence-based quantification of tumor progression showed that NT157 decreased total flux compared to the vehicle controls (Figures 4C and D). This suggests a tumor size decrease with NT157 treatment. These results suggest that inhibiting IRS1/2 can block UM tumor growth.

**Figure 4.**
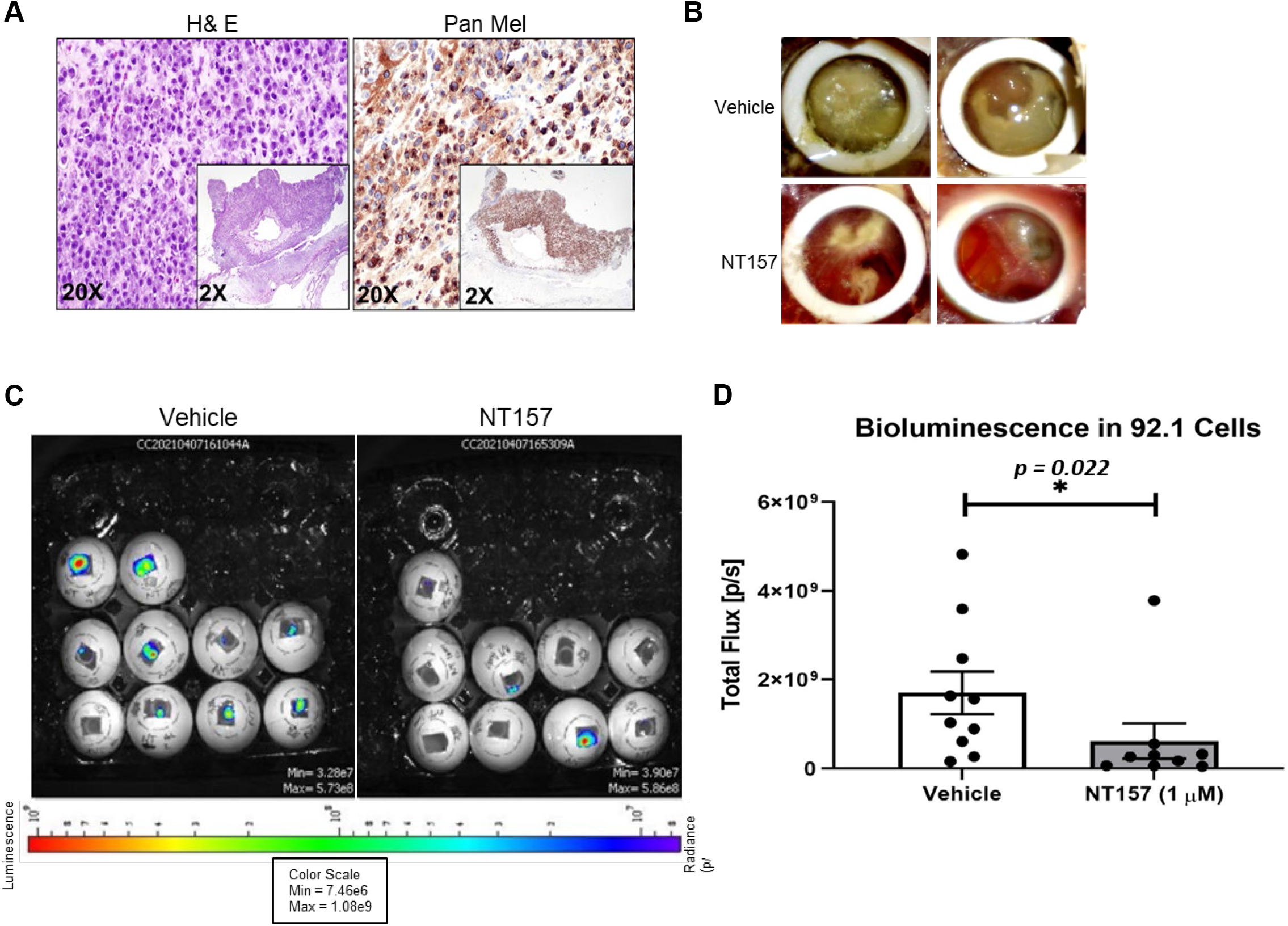
NT157 treatment reduces UM tumor growth in the chicken CAM model. (**A**) H&E (left) and pan-melanoma cocktail (anti-HMB1, Tyrosinase and S100 antibodies; right) staining of UM tumor tissues from 92.1 cells grown in chicken CAM model show the presence of melanoma cells. Representative (20x) and high magnification (inlay 2x) images shown as described. (**B**) Brightfield images of vehicle-treated (upper panel) and NT157-treated (1 μM; lower panel) representative tumors (two representative images per treatment) show a reduction of tumor size with NT157 treatment. (**C**) Representative bioluminescence images of chicken eggs bearing luciferase-tagged UM cells (92.1) treated with vehicle controls (left panel) or NT157 (1 μM) (right panel). (**D**) Tumor size of the implants was calculated as the total flux (photons per second) from the images in (C), which shows a reduction in tumor size with NT157 treatment; mean ± SD of three independent experiments was plotted as a graph; *\*P* = 0.022.

### Inhibition of IRS-1 significantly reduces UM tumor growth in mice

To complement the chicken CAM model data, the efficacy of targeting IRS-1 using NT157 was also examined using subcutaneous UM models in mice (see Materials and Methods). Xenograft tumors were generated using the UM cell lines 92.1 and MM28 in NSG mice. Five mice were treated with vehicle and five others with NT157. Treatment with NT157 strongly suppressed UM tumor growth as compared to the untreated control models (Figures 5A and C). Significant differences between the NT157 and vehicle-treated groups were also observed in the average UM xenograft tumor weight at the end of the experiments (Figure 5B). Therefore, in these experiments, we validated *in vivo* the inhibitory effect of IRS-1 blocking on tumor cell growth. There were no significant changes in the average body weights of mice in the NT157-treated vs. vehicle-treated group demonstrating that the dose of NT157 chosen for treatment was well tolerated.

**Figure 5.**
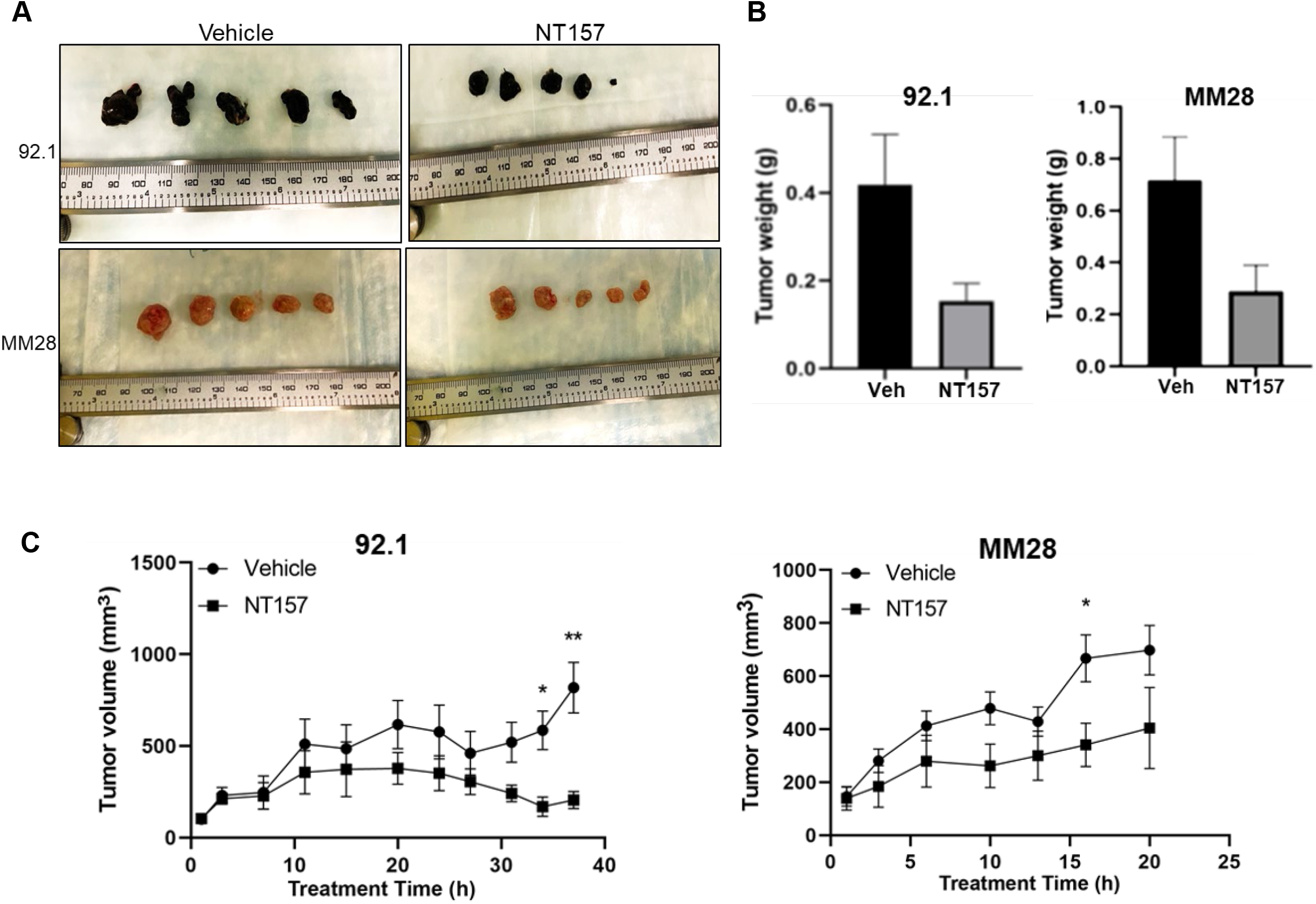
NT157 treatment inhibits UM tumor growth in a SubQ mouse model. Subcutaneous tumors grown in NSG mice with 0.5 million UM cells 92.1 (top) and MM28 (bottom) were treated intraperitoneally with NT157 (50 mg/kg body weight), 3 times per week after tumors reached ~100 mm size. Tumor volume was measured twice weekly with slide calipers. When more than 2 tumors reached 1000 mm^3^ volume in any one group, the experiment was ended and tumors were harvested. (**A**) Shows harvested tumors from untreated and NT157 treated groups at the endpoint. (**B**) Shows a reduction in collective tumor weight at the endpoint and (**C**) time-depedent reduction in tumor volume with NT157 treatement (1 μM) vs. untreated controls. The data points marked ** and *show *P* < 0.01 and <0.05, respectively.

## Discussion

Previously, we targeted IGF-1R in UM in a phase II trial with IMC-A12, a human monoclonal antibody to the receptor [11], and we did not observe any responders and concluded that this monoclonal antibody was not a viable treatment option. However, our *in vitro* experiments with this antibody showed inhibition of IGF-1R activation and cell migration [11]. In the clinical setting, monoclonal antibodies may not affect functional aspects of cell survival efficiently [32], because antibody uptake by the tumor depends on the balance between pharmacokinetics and efficient penetration as well as retention in the targeted tissue. Therefore, here, we evaluated NT157, a small molecule inhibitor of IRS-1/2, the substrates immediately downstream to IGF-1R, in this study.

After confirming IGF-1R expression in UM tumors from our preliminary TCGA analysis (Figure 1A), we also observed IGF-1R over-expression in all UM cell lines tested (Figure 1B). Higher expression of IGF-1R in the UM cells compared to non-cancerous cells (Figure 1D) shows that targetting IGF-1R in patients may circumvent toxicity challenges. IRS-1 inhibition, using NT157, successfully blocked UM cell survival, cell migration and induced apoptosis in UM cells (Figure 2). Our RPPA analysis, post -NT157 treatment, identified key signaling pathways/proteins that were altered upon IRS-1 inhibition in UM (Figure 3). Most importantly, we observed an increase in protein levels of p85 subunit of PI3K, indicating lower PI3K activation upon NT157 treatment. Since PI3K/AKT pathway promotes cell survival, reduced activation of this pathway might contribute to tumor growth inhibition, as evident from our *in vivo* data (Figures 4 and 5). We also observed changes in protein levels of other cell signaling pathways (hexokinase-II, MAPK, ribosomal protein S6 [RPS6], and STAT3) upon NT157 treatment (Figure 3A), however these genes were not associated with overall UM survival (data not shown) when we analyzed the TCGA data sets [33]. For these markers, we compared low (below median expression) and high (above median) groups and found no statistically significant associations.

We also compared a monoclonal neutralizing antibody of IGF-1R against NT157 in their capacity to block UM cell survival and UM cell migration (Figure S1). Cancer cell migration is an important step for disease progression and metastasis. Since, UM with primary tumor in the eye shows distant metastasis to liver (>90%) [2, 3], liver-borne IGF-1 may play a major role in UM cell migration. We have shown that NT157 performs better than the monoclonal antibody in each of these assays and is therefore, a better candidate for inhibiting IGF-1/IGF-1R signaling in UM cell models (Figure S1). The success of a therapeutic strategy involving a small molecule, or an antibody depends on multiple factors like uptake, bioavailability, and stability of the agent under physiological conditions. Therefore, it is important to test both these avenues, pre-clinically, to develop the most effective or suitable therapeutic strategy.

Most importantly, we validated the *in vitro* inhibitory effect of NT157 on UM cell survival using *in vivo* models for UM tumor growth. Tumor growth reduction in a chicken CAM model of UM with NT157 treatment clearly shows that blocking the IGF-1/IGF-1R axis in UM is beneficial in controlling tumor growth (Figure 4). Similarly, in our subQ mouse model of UM tumor growth, NT157 treatment resulted in reduction of tumor volume and tumor weight (Figure 5). Since NT157 is not a clinical compound, our study forms the basis or proof-of-concept for targeting IGF-1/IGF-1R in a clinical setting in UM.

In conclusion, our results suggest blocking IGF-1/IGF-1R axis in UM inhibits cell survival, cell migration, and induces apoptosis *in vitro*. Targeting this pathway *in vivo* inhibits UM tumor growth. Recent FDA approval of Tebentafusp for use in non-resectable UM [34, 35] has generated hope and created a therapeutic option for a section of the metastatic UM patients. Since this is the only approved therapy and one size does not fit all (tebentafusp only is applicable to HLA-A*02:01-positive adult patients), there is an urgent need to develop new therapeutic strategies. Our results strongly support the rationale to develop clinical studies based on IGF-1/IGF-1R inhibition in UM. Considering the fact that single-agent therapies often do not produce durable responses in melanoma, the future therapeutic strategies blocking IGF-1/IGF-1R axis may need to consider combination therapies for a synergistic effect to control tumor growth.

## Author contributions

Concept and design: CC, EAG; Acquisition of data: CC, Rajat Bhattacharya, YQ, FSK, GAW, HV, and Rishav Bhattacharya; Chicken CAM model-based study design and execution: GAW and HV; High throughput drug screening and analyses: CS and PR, respectively; Analysis and interpretation of data: CC, JR, SP, and HV; Writing, review, and/or revision of the manuscript: CC, JR, SP, HV, and EAG; Study supervision: CC

## Acknowledgements

We thank Dr. Sirisha Yadugiri for manuscript reviewing and editing, and Mrs. Violet Garcia for manuscript formatting and administrative support. We thank Dr. Michael A. Davies for scientific advice and critical review to improve the quality of the manuscript. We also thank The UT MD Anderson Flow Cytometry and cellular imaging facility, which is supported in part by NIH through the MD Anderson Cancer Center support grant CA016672, for providing support for FACS analysis and imaging. We thank The Patient-Derived Xenograft and Advanced in vivo Models Core Facility, which is supported in part by NIH through the Baylor College of Medicine Cancer Center support grant #2P30CA125123-14. We also thank The UT MD Anderson RPPA core, supported by NCI grant # CA16672 and NIH grant R50CA221675, for RPPA data analysis.

## Data Availability Statement

All relevant data are available from the corresponding author upon request.

## Supplemental Figures

**Supplemental Figure 1.**
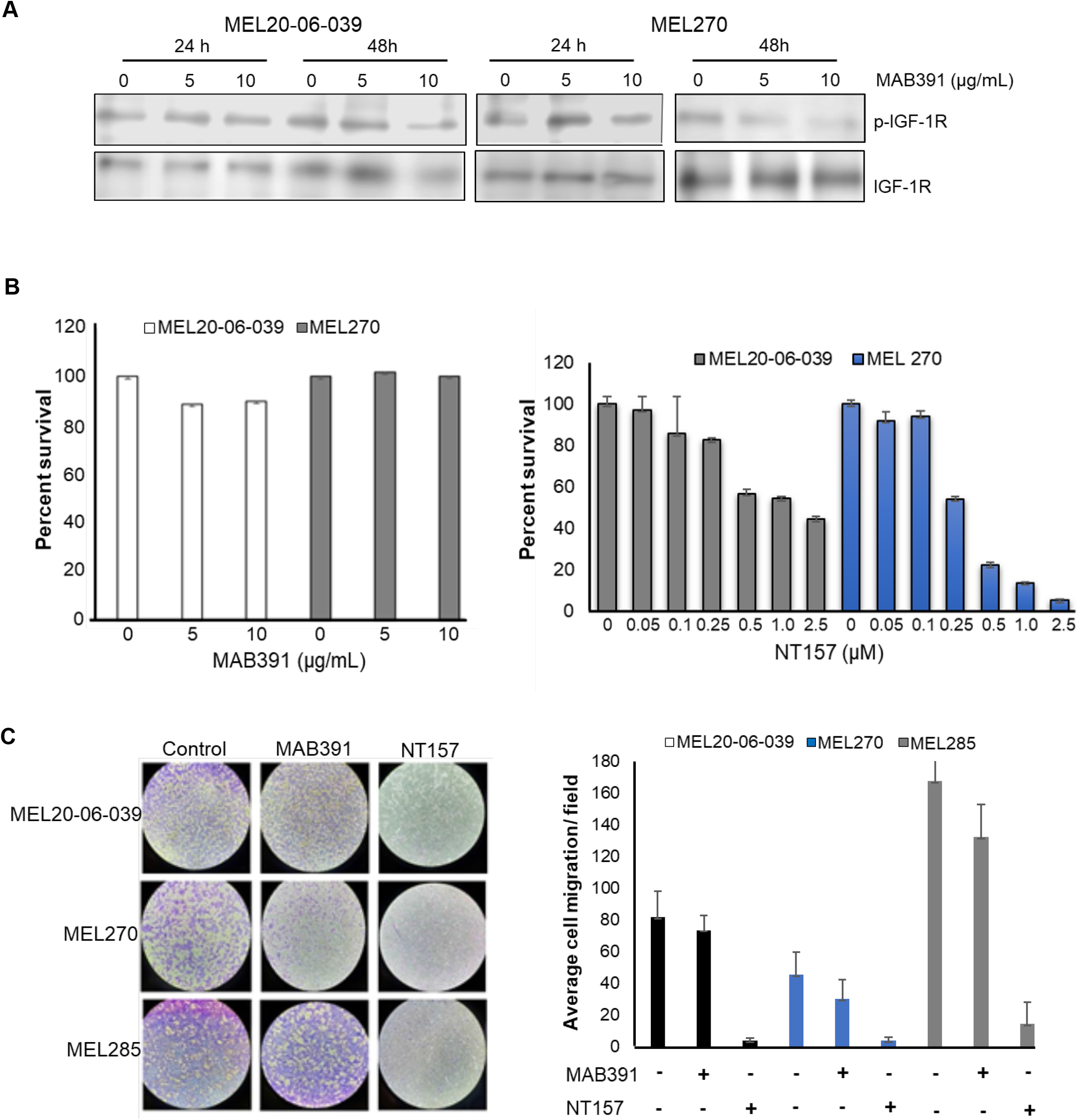
NT157 is more efficient in inhibiting UM cell survival and migration compared to anti-IGF-1R antibody. (**A**) Western blot analysis of IGF-1R activation with MAB391 in UM cells (MEL20-06-039 and MEL270 cell lines) shows dose and time-dependent inhibition of IGF-1R phosphorylation, confirming the specificity of the antibody. (**B**) Quantification of cell survival measured through MTT-based assay after treatment of UM cells (MEL20-06-039 and MEL270 cell lines) with MAB391(left) or NT157 (right) or vehicle control, shows NT157 is more effective in blocking UM cell survival compared to MAB391. The bar graphs represent percent survival and mean ± SD of three independent experiments were plotted. (**C**) An *in vitro* cell migration assay shows the staining of migrated UM cells (MEL20-06-039, MEL270, MEL285 cell lines; left). The quantification of stained cells (right) shows NT157 (2.5 μM) treatment blocks cell migration more efficiently than MAB391 (100 ng/mL) or vehicle controls. Bar graphs represent the mean ± SD of average number of cells counted/field from 5 fields per condition, in two independent set of experiments.

**Supplemental Figure 2.**
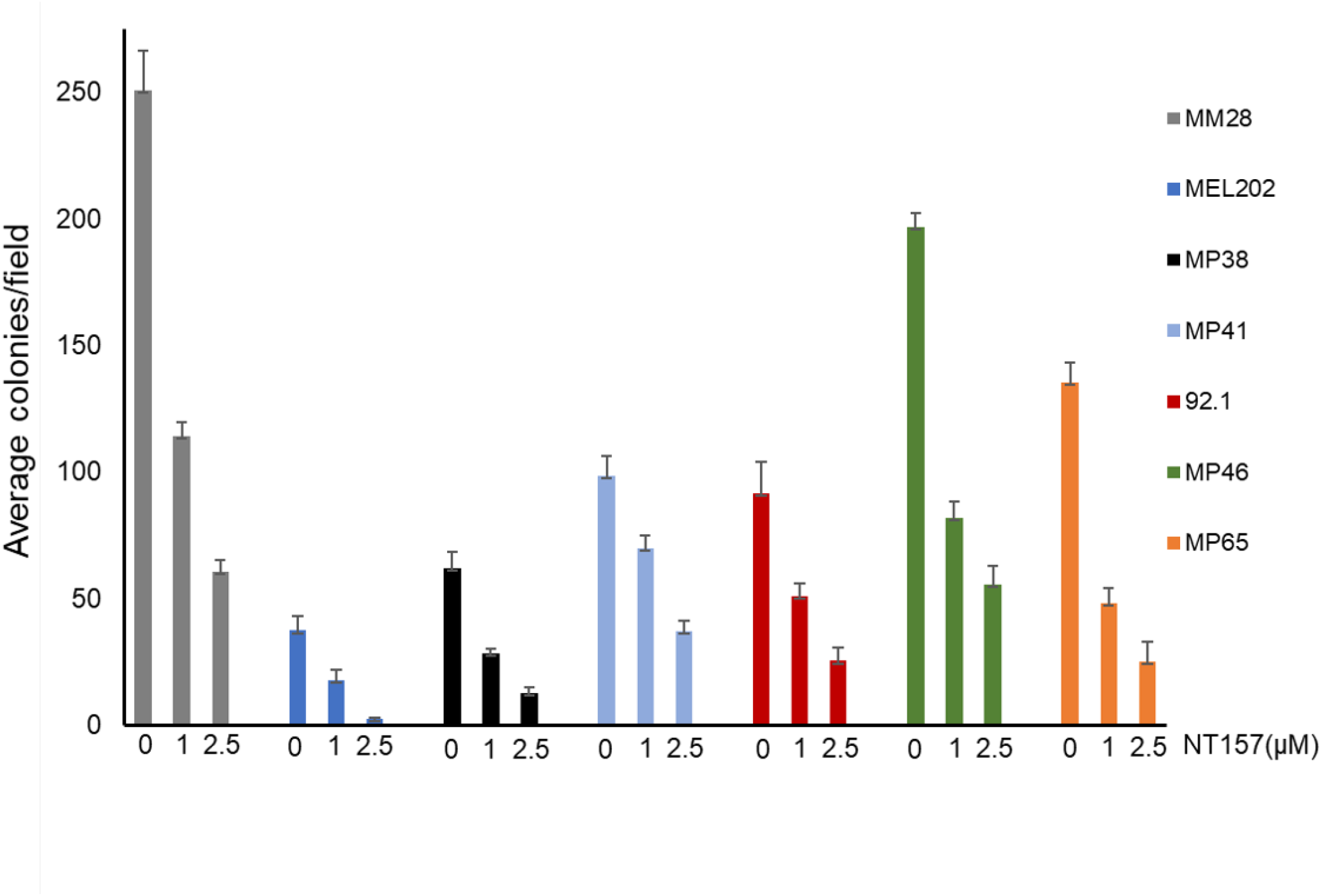
Effect of NT157 on UM cell survival in a colony formation assay. The average number of colonies counted after staining with crystal violet in NT157 treated (1 and 2.5 μM) and untreated control UM cell lines (MM28, MEL202, MP38, MP41, 92.1, MP46, MP65) were plotted, which shows dose-dependent inhibition of UM cell survival with NT157 treatment. A total of 5 fields/condition/cell line were counted to obtain the average number of colonies, in two independent set of experiments.

